## Supplementary Figures for "Nageotte nodules in human DRG reveal neurodegeneration in painful diabetic neuropathy"

**List of Supplemental Materials:**
Figs. S1 to S11
Tables S1 to S6
Movies S1 to S3

### Supplementary Figures

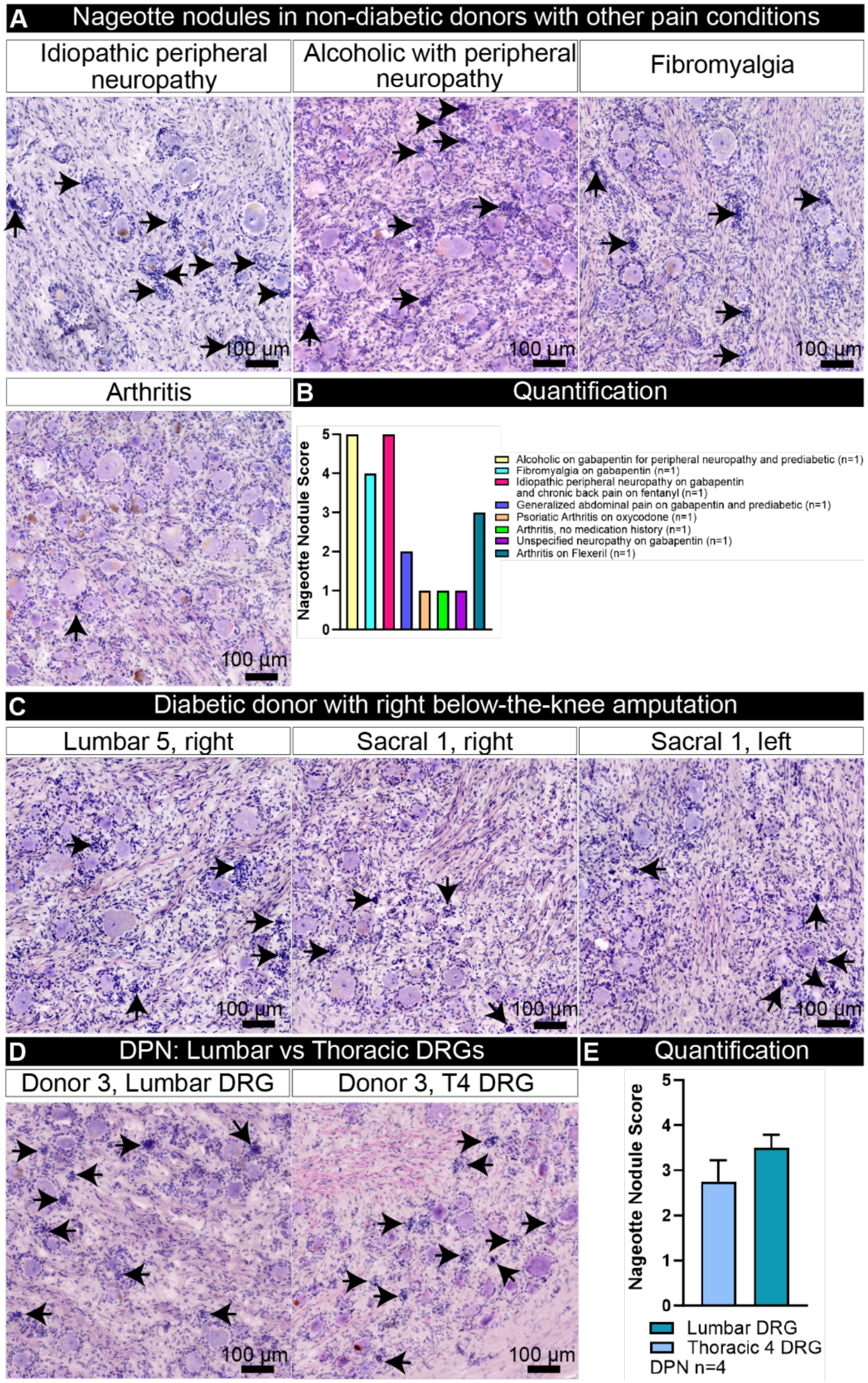


**Supplemental Figure 1. Nageotte nodules in DRGs from organ donors with other pain conditions. A)** Images of Hematoxylin and Eosin stained DRGs from a subset of organ donors with other pain conditions**.** Black arrows point towards Nageotte nodules. **B)** Nageotte nodules were found at higher levels in donors with neuropathic conditions, but at lower levels in organ donors with a history of arthritis. **C)** Hematoxylin and eosin staining of DRGs procured from a diabetic donor with a right below-the-knee amputation. Nageotte nodules were found in DRGs that innervate the leg and foot (lumbar 5 right, sacral 1 right, and sacral 1 left) on both sides of the body. Black arrows point towards Nageotte nodules. **D)** Hematoxylin and eosin staining of a lumbar and thoracic 4 (T4) DRG from a DPN donor (donor 3). Black arrows point towards Nageotte nodules. **E)** DPN lumbar and T4 DRGs had similar Nageotte nodule scores. **Scale bars:** A, C, D: 100 μm.

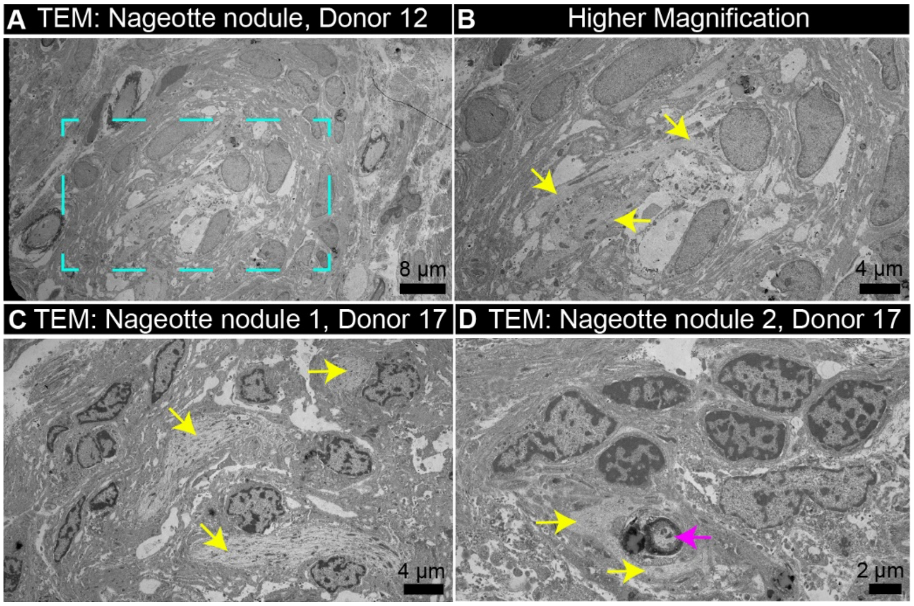


**Supplemental Figure 2. Transmission electron microscopy of Nageotte nodules. A)** Transmission electron microscopy (TEM) image of a Nageotte nodule from Donor 12. Cyan box outlines an **B)** area of the Nageotte nodule at higher magnification. Yellow arrows point to unmyelinated axonal fibers interspersed with nuclei of non-neuronal cells. **C)** TEM image of a Nageotte nodule from Donor 17 with unmyelinated fibers (yellow arrows) intermixed with non-neuronal cells. **D)** TEM image of another Nageotte nodule from Donor 17 with unmyelinated fibers (yellow arrows) and a slightly myelinated fiber (magenta arrow) intermixed with the nuclei of other cells. **Scale bars:** A: 8 μm. B-C: 4 μm. D: 2 μm.

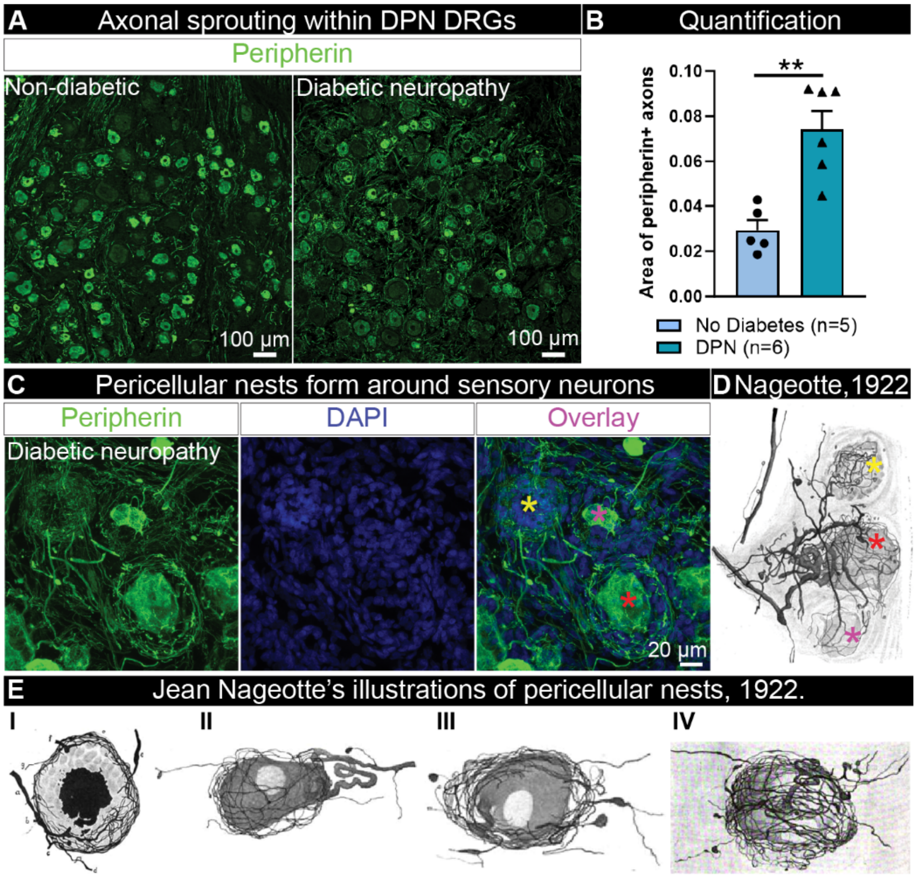


**Supplemental Figure 3. Peripherin-positive axonal fibers sprout throughout the DPN DRGs, forming Nageotte nodule axon bundles and pericellular nests. A)** Representative 20X confocal images of a non-diabetic and diabetic peripheral neuropathy (DPN) DRG labeled for peripherin (green). Axonal sprouting was observed throughout the DPN DRGs, and when quantified, **B)** the area of peripherin-positive axons within the neuron-rich area of the DRG was significantly elevated in the DPN DRGs compared to non-diabetic controls. **C)** 60X projected z-stack image of peripherin-positive fibers intertwining at a Nageotte nodule (yellow asterisk) and forming pericellular nests (PCNs) around two sensory neurons, one with a visible cell body (red asterisk) and another with a misshapen/shrunken soma (magenta asterisk) possibly in a state of dying. **D)** Jean Nageotte's illustration from 1922 of a similar morphology showing a Nageotte nodule (yellow asterisk), a sensory neuron ensheathed by a PCN formed by its own neurites, and a second dying neuron (magenta asterisk) that is ensheathed by a PCN. The axons at the Nageotte nodule and the PCN around the dying neuron (magenta asterisk) are formed by fibers sprouting from the glomerulus of the intact neuron (red asterisk). **E)** Jean Nageotte's other illustrations of PCNs. I) a dead neuron with delayed phagocytosis. II) PCN formed by neurites coming from the same neuron. III) PCN formed by branches born at the end of the surviving portion of the glomerulus. IV) PCN formed by branches from the glomerulus. **Scale bars:** A: 100 µm. C: 20 µm. **Sample size:** Non-diabetic n=5, DPN n=6.

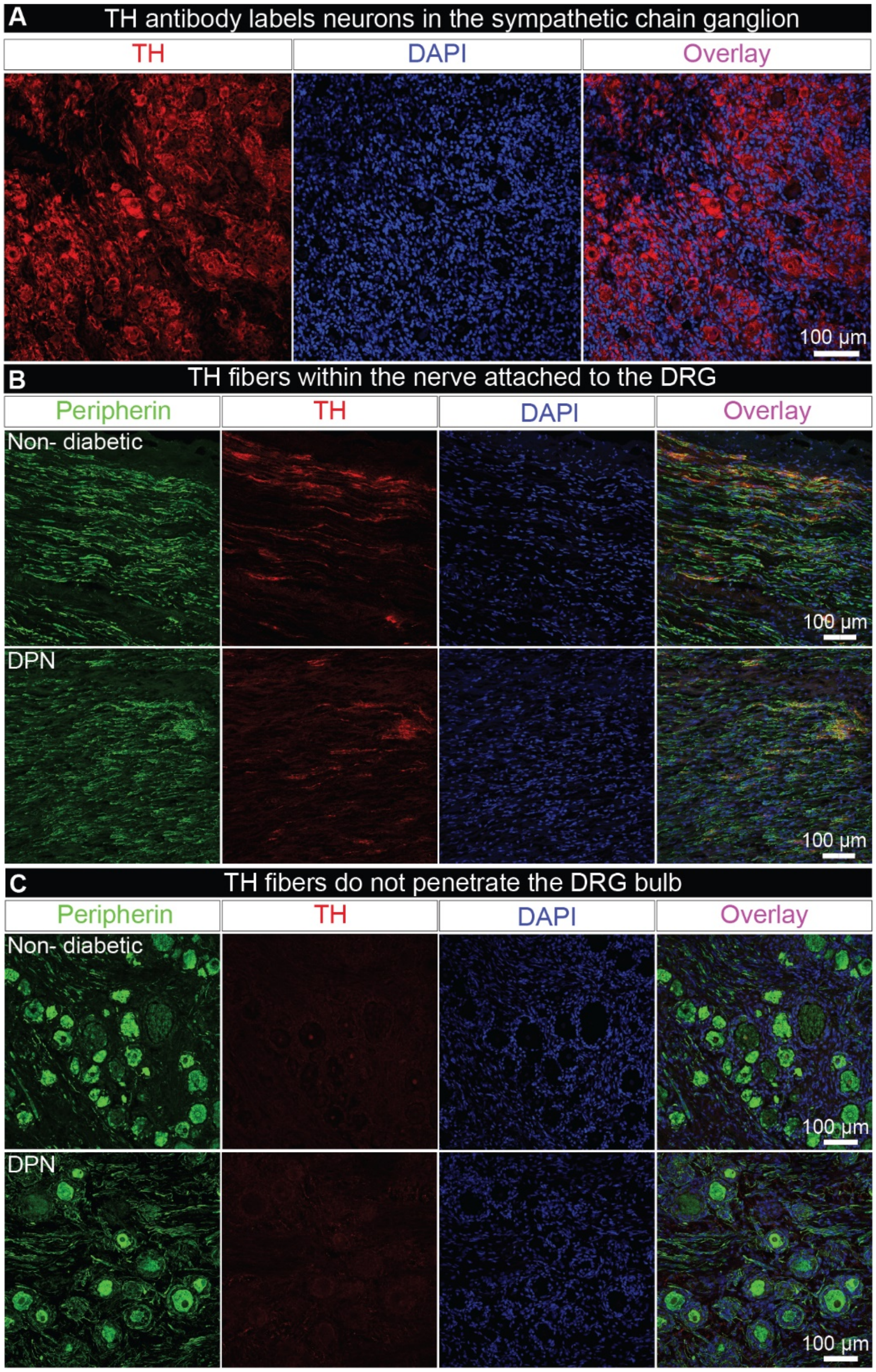


**Supplemental Figure 4. Tyrosine Hydroxylase staining in non-diabetic and DPN DRGs. A)** The tyrosine hydroxylase (TH) antibody (ab152, red) robustly labels sympathetic neurons in the human sympathetic chain ganglion. **B)** Representative 20X confocal images of the nerve region of a non-diabetic and diabetic peripheral neuropathy (DPN) DRG labeled for peripherin (green), TH (red), and DAPI (blue). Sparse TH+ fibers were detected in the nerve attached to the DRG. **C)** Representative 20X confocal images of the neuron-rich region of a non-diabetic and DPN DRG labeled for peripherin (green), TH (red), and DAPI (blue). Little-to-no TH+ fibers were detected in the neuron-rich area of the DRG. **Scale bars:** A: 50 µm. B-C: 100 µm. **Sample size:** Non-diabetic n=6, DPN n=6.

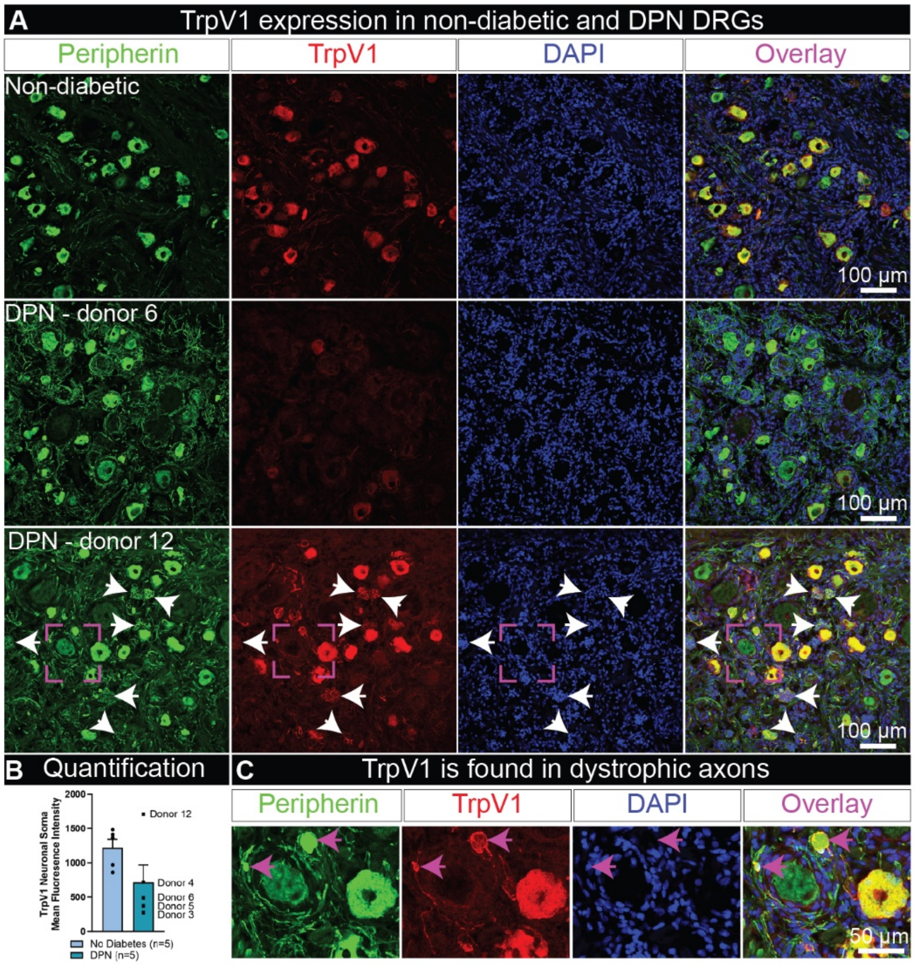


**Supplemental Figure 5. TrpV1 expression in diabetic painful neuropathy DRGs. A)** Representative 20X confocal images of peripherin (green), TrpV1 (red), and DAPI (blue) in a non-diabetic DRG, and two DPN DRGs. Donor 6 had a medical history indicating neuropathy of the feet, and Donor 12 was taking analgesics, had difficulty walking but had no established diagnosis of neuropathy. White arrows point towards Trpv1+ Nageotte nodule axonal bundles. **B)** Quantification of the mean fluorescence intensity of TrpV1 in the neuronal soma of all sensory neurons. Donor 12 had elevated TrpV1 expression, while the other DPN donors with medical notes of neuropathy or diabetes-related amputation showed reduced Trpv1 expression within the DRGs. **C)** In Donor 12, TrpV1 was also detected in dystrophic axons (magenta arrows). **Scale bars:** A: 100 µm. C: 50 µm. **Statistical test**: unpaired t-test **Sample size:** Non-diabetic n=5, DPN n=5.

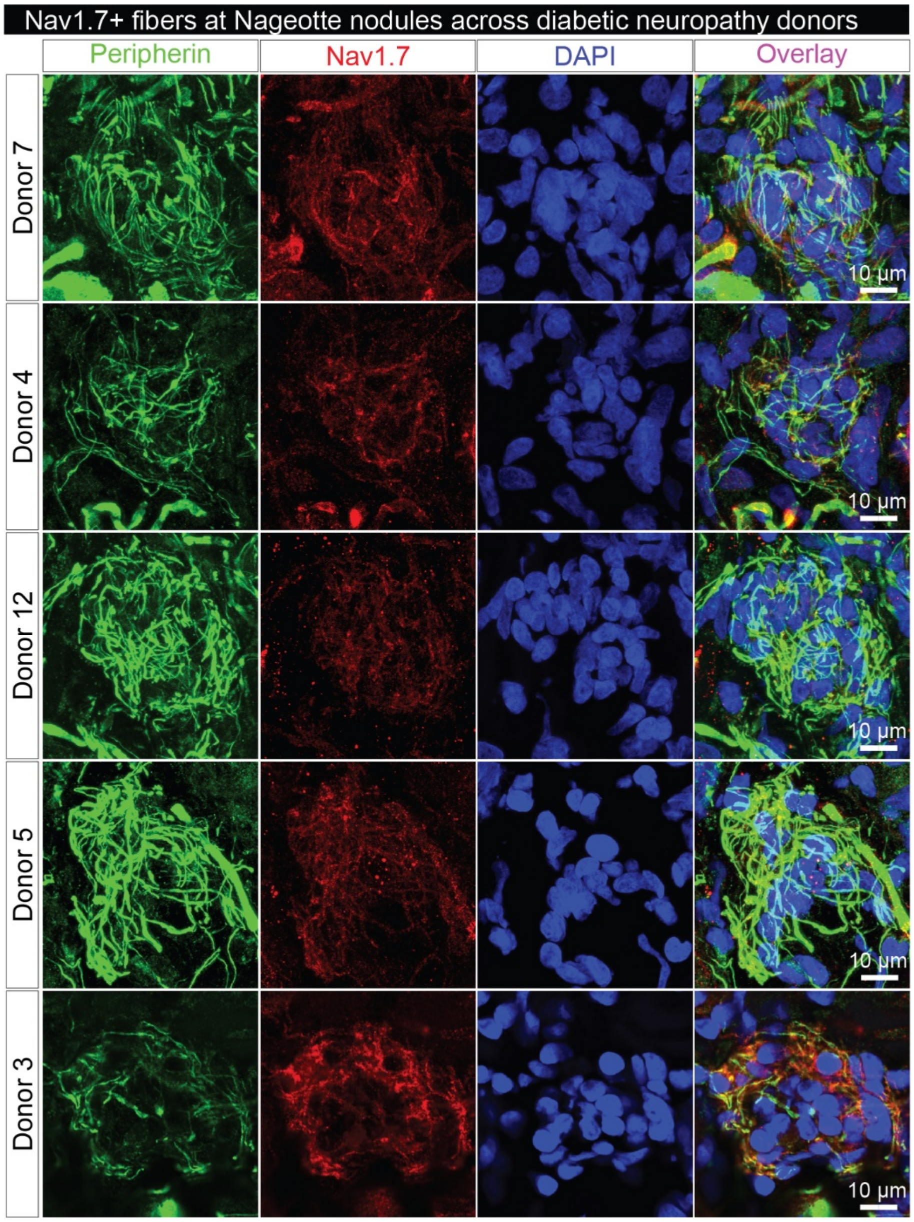


**Supplemental Figure 6. Nav1.7 expression in Nageotte nodules in diabetic peripheral neuropathy DRGs.** Representative 100X confocal images of peripherin (green), Nav1.7 (red), and DAPI (blue) in a Nageotte nodule from DPN DRGs. **Scale bars:** 10 µm. **Sample size:** DPN n=5.

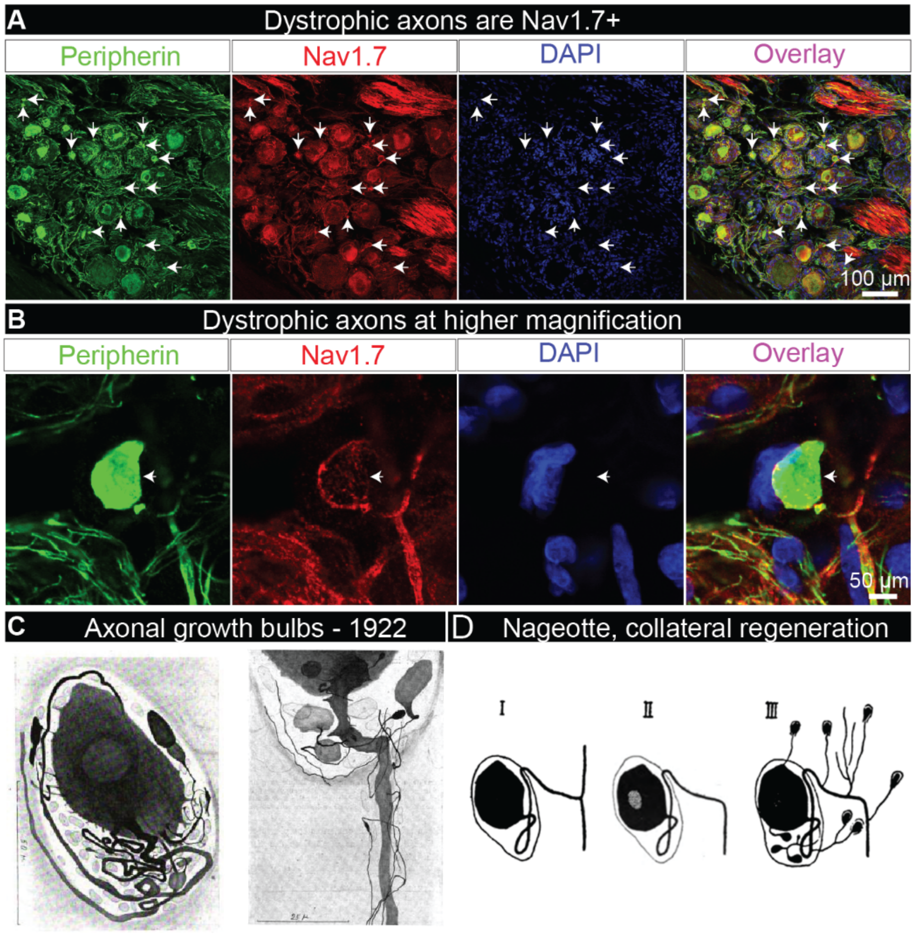


**Supplemental Figure 7. Nav1.7 expression in dystrophic axons in diabetic painful neuropathy DRGs. A)** Representative 20x confocal images of a DPN DRG labeled for peripherin (green), Nav1.7 (red), and DAPI (blue). Nav1.7 was robustly detected in axonal fibers within the DRG including dystrophic axons (white arrows) **B)** Representative 100X confocal images of a dystrophic axon (white arrow) in a DPN DRG labeled for peripherin (green), Nav1.7 (red), and DAPI (blue) **C)** Jean Nageotte described dystrophic axons in 1922 as "growth balls" which he claimed arose from the sensory neuron soma, glomerulus, and extracapsular portion of the axon in a process known as **D)** collateral regeneration. C and D are original figures from Jean Nageotte, 1922. **Scale bars:** A: 100 µm. B: 5 µm. Sample size: DPN n=5.

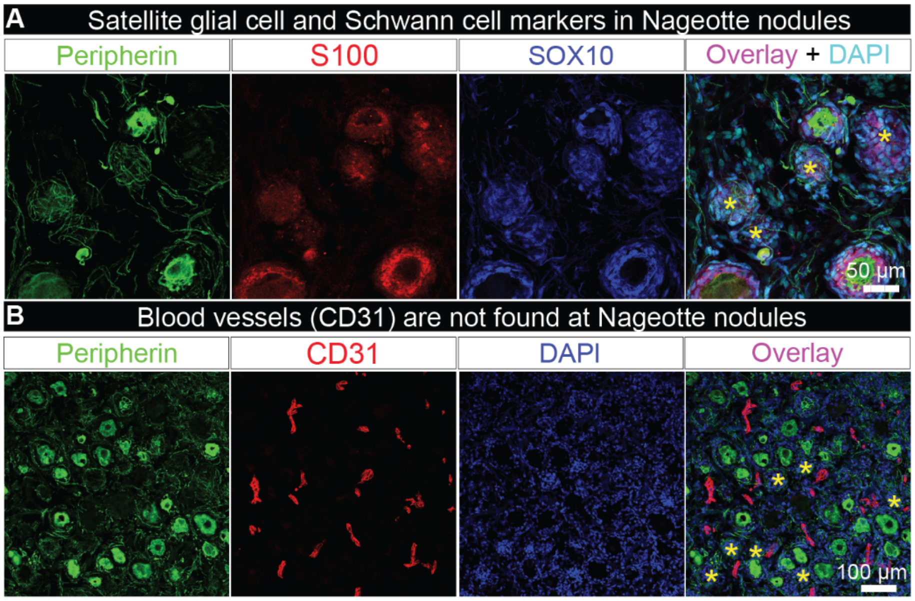


**Supplemental Figure 8. Immunohistochemistry for markers of satellite glial cells, Schwann cells, and blood vessels in DPN DRGs. A)** Representative 40X confocal image of a DPN DRG labeled for peripherin (green), S100 (red), and SOX10 (blue), and DAPI (cyan). S100 and SOX10 label satellite glia and Schwann cells and were detected at Nageotte nodules. Sample size: DPN n=6. B) Representative 20X confocal image of a DPN DRG labeled for peripherin (green), CD31 (red), and DAPI (blue). CD31, also known as PECAM-1, is a platelet/endothelial cell adhesion molecule that labels blood vessels. CD31 was not detected in Nageotte nodules. Sample size: DPN n=5. Scale bars: A: 50 µm. B: 100 µm.

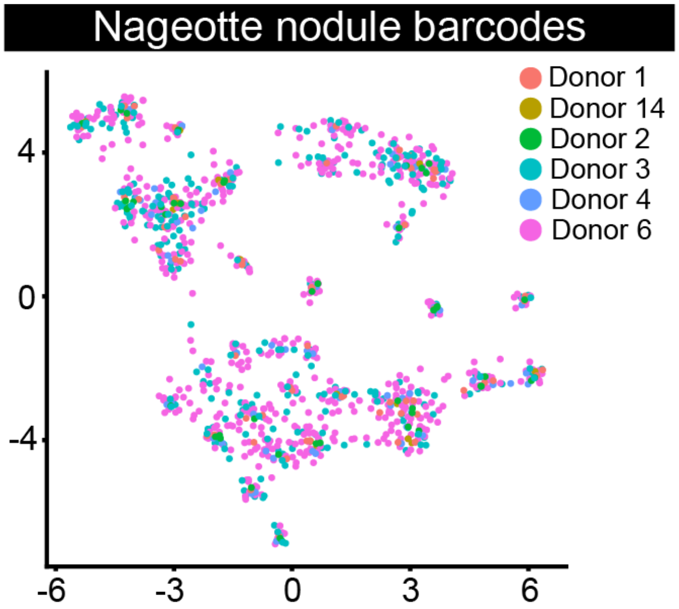


**Supplemental Figure 9. Distribution of Nageotte nodule barcodes from each donor across clusters and disease associated glia gene signature.** Nageotte nodule barcodes from each donor is represented in each cluster.

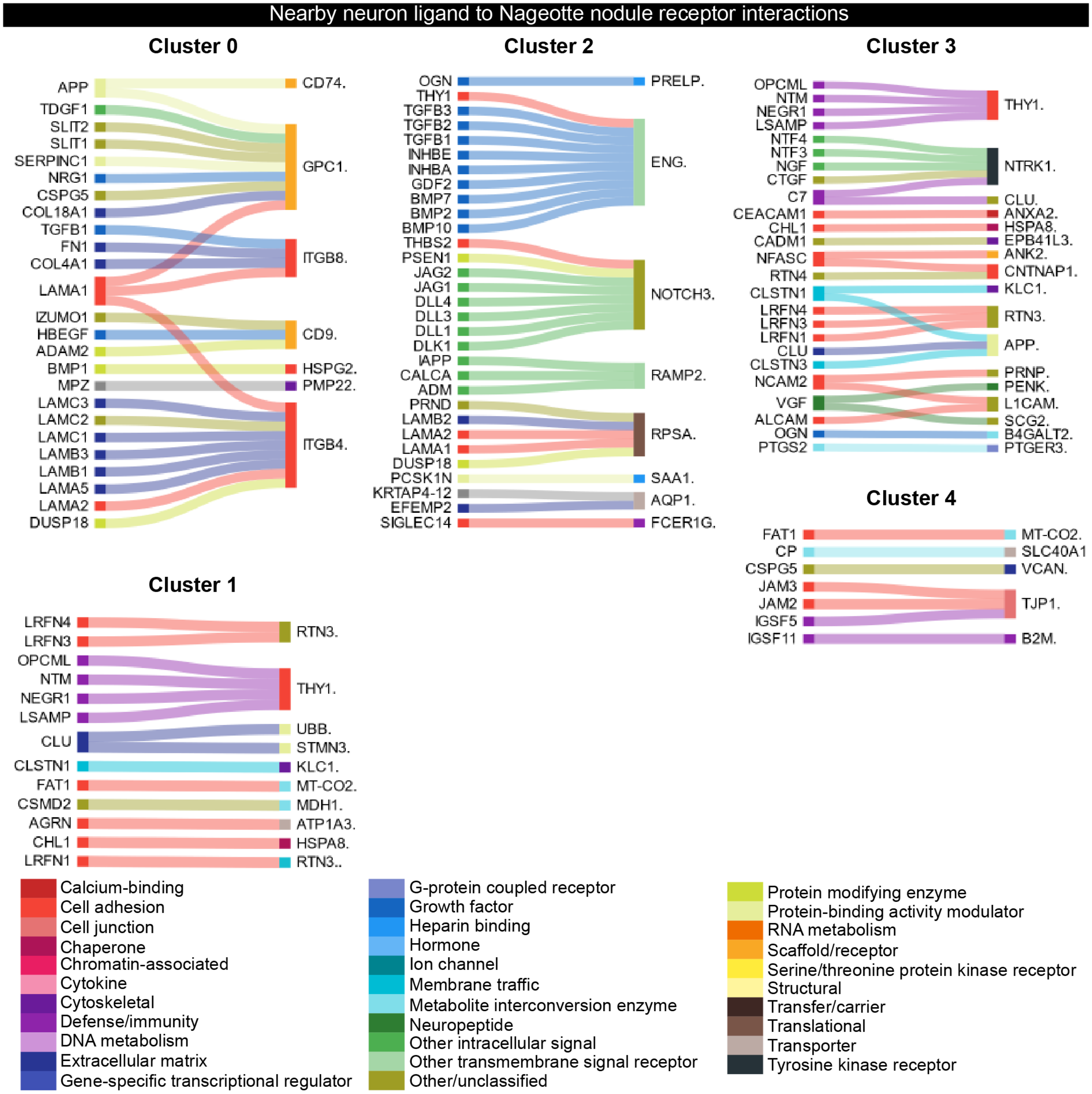


**Supplemental Figure 10. Ligand-receptor interactions between nearby neurons and Nageotte nodules.** Differentially expressed receptors per Nageotte nodule cluster and corresponding ligands expressed in nearby neurons.

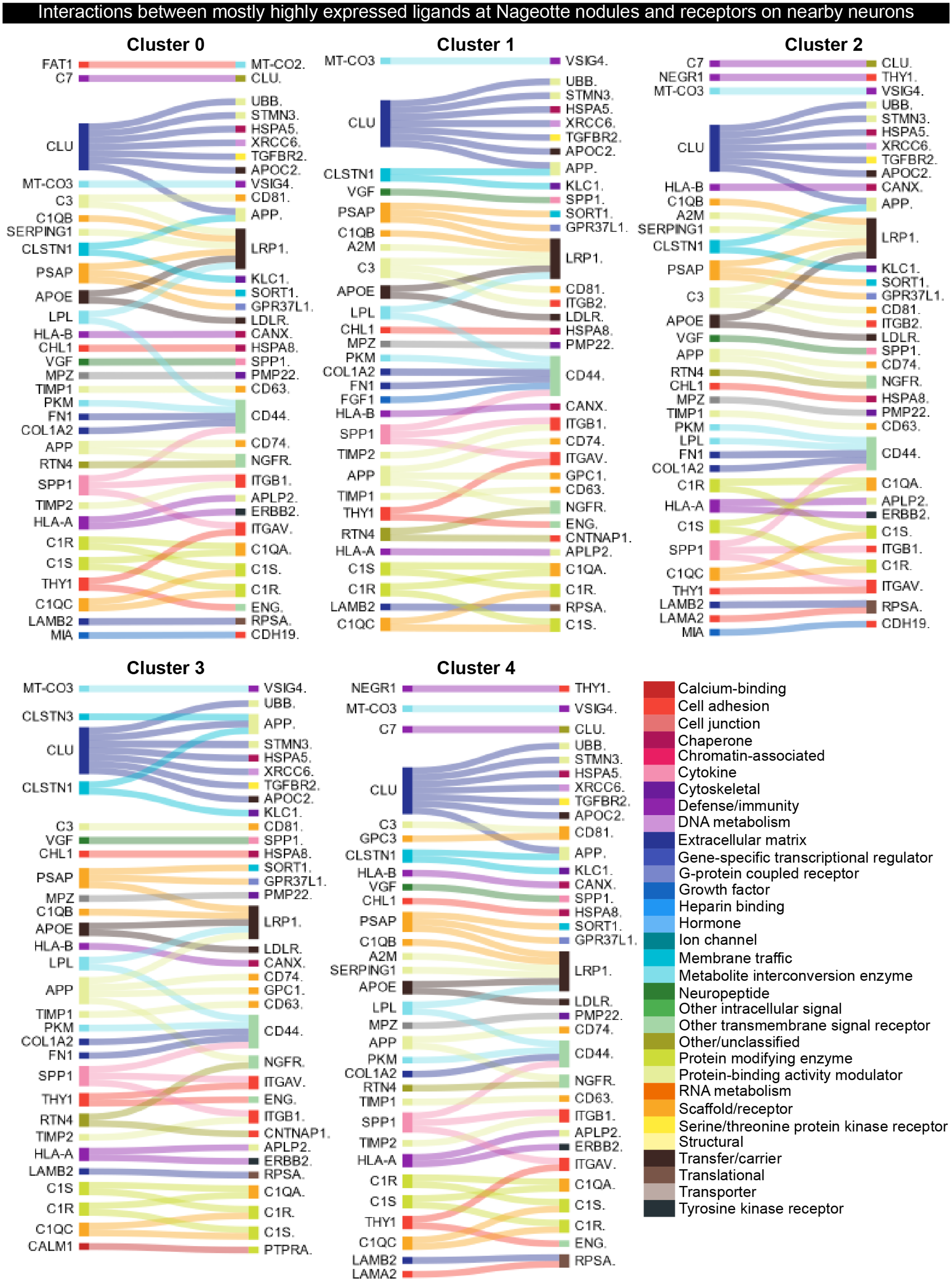

Interactions between mostly highly expressed ligands at Nageotte nodules and receptors on nearby neurons
Cluster 0
FAT1
C7
CLU
MT-CO3
C3
C1QB
SERPING1
CLSTN1
PSAP
APOE
LPL
HLA-B
CHL1
VGF
MPZ
TIMP1
PKM
FN1
COL1A2
APP
RTN4
SPP1
TIMP2
HLA-A
C1R
C1S
THY1
C1QC
LAMB2
MIA
MT-CO2.
CLU.
UBB.
STMN3.
HSPA5.
XRCC6.
TGFBR2.
APOC2.
VSIG4.
CD81.
APP.
LRP1.
KLC1.
SORT1.
GPR37L1.
LDLR.
CANX.
HSPA8.
SPP1.
PMP22.
CD63.
CD44.
CD74.
NGFR.
ITGB1.
APLP2.
ERBB2.
ITGAV.
C1QA.
C1S.
C1R.
ENG.
RPSA.
CDH19.
Cluster 1
MT-CO3
CLU
CLSTN1
VGF
PSAP
C1QB
A2M
C3
APOE
LPL
CHL1
MPZ
PKM
COL1A2
FN1
FGF1
HLA-B
SPP1
TIMP2
APP
TIMP1
THY1
RTN4
HLA-A
C1S
C1R
LAMB2
C1QC
VSIG4.
UBB.
STMN3.
HSPA5.
XRCC6.
TGFBR2.
APOC2.
APP.
KLC1.
SPP1.
SORT1.
GPR37L1.
LRP1.
CD81.
ITGB2.
LDLR.
HSPA8.
PMP22.
CD44.
CANX.
ITGB1.
CD74.
ITGAV.
GPC1.
CD63.
NGFR.
ENG.
CNTNAP1.
APLP2.
C1QA.
C1R.
RPSA.
C1S.
Cluster 2
C7
NEGR1
MT-CO3
CLU
HLA-B
C1QB
A2M
SERPING1
CLSTN1
PSAP
C3
APOE
VGF
APP
RTN4
CHL1
MPZ
TIMP1
PKM
LPL
FN1
COL1A2
C1R
HLA-A
C1S
SPP1
C1QC
THY1
LAMB2
LAMA2
MIA
CLU.
THY1.
VSIG4.
UBB.
STMN3.
HSPA5.
XRCC6.
TGFBR2.
APOC2.
CANX.
APP.
LRP1.
KLC1.
SORT1.
GPR37L1.
CD81.
ITGB2.
LDLR.
SPP1.
CD74.
NGFR.
HSPA8.
PMP22.
CD63.
CD44.
C1QA.
APLP2.
ERBB2.
C1S.
ITGB1.
C1R.
ITGAV.
RPSA.
CDH19.
Cluster 3
MT-CO3
CLSTN3
CLU
CLSTN1
C3
VGF
CHL1
PSAP
MPZ
C1QB
APOE
HLA-B
LPL
APP
TIMP1
PKM
COL1A2
FN1
SPP1
THY1
RTN4
TIMP2
HLA-A
LAMB2
C1S
C1R
C1QC
CALM1
VSIG4.
UBB.
APP.
STMN3.
HSPA5.
XRCC6.
TGFBR2.
APOC2.
KLC1.
CD81.
SPP1.
HSPA8.
SORT1.
GPR37L1.
PMP22.
LRP1.
LDLR.
CANX.
CD74.
GPC1.
CD63.
CD44.
NGFR.
ITGAV.
ENG.
ITGB1.
CNTNAP1.
APLP2.
ERBB2.
RPSA.
C1QA.
C1R.
C1S.
PTPRA.
Cluster 4
NEGR1
MT-CO3
C7
CLU
C3
GPC3
CLSTN1
HLA-B
VGF
CHL1
PSAP
C1QB
A2M
SERPING1
APOE
LPL
MPZ
APP
PKM
COL1A2
RTN4
TIMP1
SPP1
TIMP2
HLA-A
C1R
C1S
THY1
C1QC
LAMB2
LAMA2
THY1.
VSIG4.
CLU.
UBB.
STMN3.
HSPA5.
XRCC6.
TGFBR2.
APOC2.
CD81.
APP.
KLC1.
CANX.
SPP1.
HSPA8.
SORT1.
GPR37L1.
LRP1.
LDLR.
PMP22.
CD74.
CD44.
NGFR.
CD63.
ITGB1.
APLP2.
ERBB2.
ITGAV.
C1QA.
C1S.
C1R.
ENG.
RPSA.
Calcium-binding
Cell adhesion
Cell junction
Chaperone
Chromatin-associated
Cytokine
Cytoskeletal
Defense/immunity
DNA metabolism
Extracellular matrix
Gene-specific transcriptional regulator
G-protein coupled receptor
Growth factor
Heparin binding
Hormone
Ion channel
Membrane traffic
Metabolite interconversion enzyme
Neuropeptide
Other intracellular signal
Other transmembrane signal receptor
Other/unclassified
Protein modifying enzyme
Protein-binding activity modulator
RNA metabolism
Scaffold/receptor
Serine/threonine protein kinase receptor
Structural
Transfer/carrier
Translational
Transporter
Tyrosine kinase receptor

**Supplemental Figure 11. Top 50 ligand-receptor interactions between nearby neurons and Nageotte nodules per Nageotte nodule cluster.** Most highly expressed ligands (ligand genes in the top 10% of genes in each Nageotte nodule cluster) per Nageotte nodule cluster and corresponding receptors expressed in nearby neurons (receptor genes in the top 10% of all expressed genes in nearby neurons).

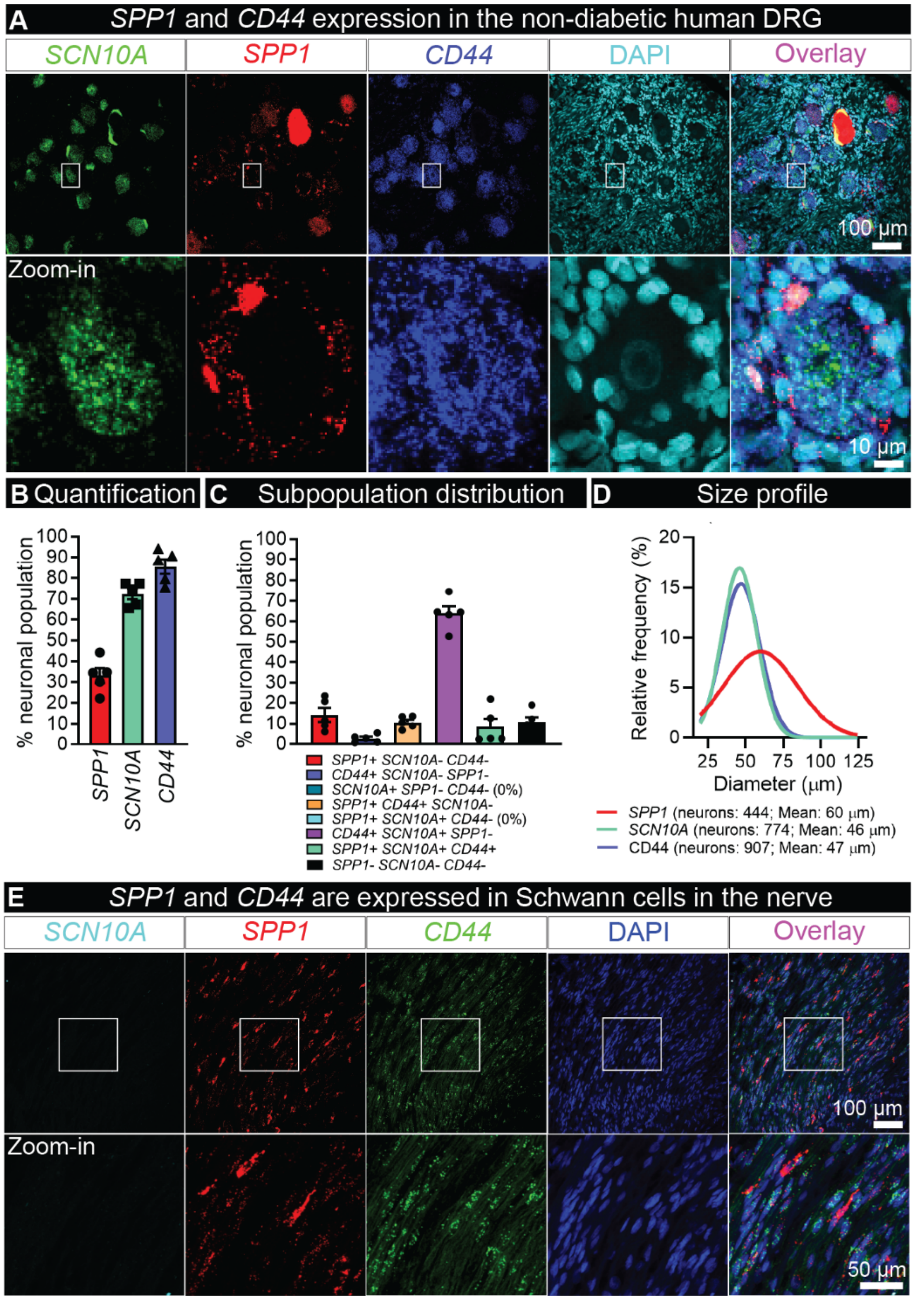


**Supplemental Figure 12. Osteopontin (*SPP1*) and CD44 mRNA expression in the non-diabetic human DRG. A)** Representative 20X confocal image of a non-diabetic DRG labeled for *SCN10A* (Nav1.8, green), *SPP1* (Osteopontin, red), and *CD44* (blue) mRNAs using RNAscope *in situ* hybridization and co-stained with DAPI (cyan). Bottom panel is a digitally magnified image of a single sensory neuron that is co-positive for *SCN10A* and *CD44*, but negative for *SPP1*. *SPP1* and CD44 were detected in the cells ringing the sensory neurons, likely satellite glial cells. **B)** Quantification of *SPP1*, *SCN10A*, and *CD44* neuronal expression in relation to the entire neuronal population within human DRG. **C)** Neuronal subpopulation distribution of *SPP1*, *SCN10A*, and *CD44*. *SPP1* was robustly detected in ~10% of the neurons that were negative for *SCN10A* and *CD44*, likely representing the proprioceptive subpopulation of sensory neurons. *SPP1* was also detected at lower levels in other subpopulations. **D)** Size profile of sensory neurons that were *SPP1*, *SCN10A*, or *CD44* positive. **E)** *SPP1* and *CD44* were also detected in cells along axonal fibers in the nerve area attached to the DRG. These cells are likely Schwann cells given their elongated nuclear shape. **Scale bars:** A: 100 µm top panel, 10 µm bottom panel. E: 100 µm top panel, 50 µm bottom panel. **Sample size:** Non-diabetic n=5.
